# Single-Cell RNA Sequencing Reveals Regulatory Mechanism for Trophoblast Cell-Fate Divergence in Human Peri-Implantation Embryo

**DOI:** 10.1101/567362

**Authors:** Bo Lv, Qin An, Qiao Zeng, Ping Lu, Xianmin Zhu, Yazhong Ji, Guoping Fan, Zhigang Xue

## Abstract

Multipotent trophoblasts undergo dynamic morphological movement and cellular differentiation after embryonic implantation to generate placenta. However, the mechanism controlling trophoblast development and differentiation during peri-implantation development remains elusive. In this study, we modeled human embryo peri-implantation development from blastocyst to early post-implantation stages by using an *in vitro* coculture system, and profiled the transcriptome of individual trophoblast cells from these embryos. We revealed the genetic networks regulating peri-implantation trophoblast development. While determining when trophoblast differentiation happens, our bioinformatic analysis identified T-box transcription factor 3 (TBX3) as a key regulator for the differentiation of cytotrophoblast into syncytiotrophoblast. The function of TBX3 in trophoblast differentiation is then validated by a loss-of-function experiment. In conclusion, our results provided a valuable resource to study the regulation of trophoblasts development and differentiation during human peri-implantation development.

## Background

The placenta is the interface between the fetal and maternal circulation, and plays an essential role in supporting fetus development and survival. Most placental functions are carried out by trophoblasts, which are derived from the trophectoderm (TE) in blastocysts [1]. After implantation, the trophectoderm becomes multipotent trophoblast stem cells, proliferate and differentiate into distinct trophoblast sublineages, including cytotrophoblast (CT), extravillous cytotrophoblast (EVT) and syncytiotrophoblast (ST) [2].

Previous anatomical studies using a relatively small number of human specimens have shown that the CT appears right after implantation, which then differentiates into the EVT and ST before day 21 post-fertilization [3]. Recent studies using single cell RNA-seq on human placenta successfully revealed transcriptomic and functional heterogeneity between different trophoblast sublineages within placenta. However, all these single cell profiling studies used placenta after 6 gestational weeks [4-7]. Due to the lacking of data for early-post implantation embryos, it is still elusive when trophoblast sublineages are established, and how the trophoblast differentiation is regulated.

Human embryos cocultured with endometrial cells *in vitro* can model morphological and molecular changes of peri-implantation embryos *in vivo* [8-10]. In this study, we profiled transcriptomes of 476 individual trophoblast cells isolated from 19 human embryos co-cultured with endometrial cells. We revealed the regulatory networks underlying trophoblast development and differentiation. Using a unique bioinformatic approach and a loss-of-function verification experiments, we identified a novel transcription factor TBX3 that regulates the trophoblast differentiation. Our results provided a rich resource to study the early placenta development.

## Results

### Single-Cell Transcriptome Profiling of Trophoblasts in Human Peri-Implantation Embryos

We first obtained human peri-implantation embryos by coculturing blastocysts with human primary endometrial cells (EM) (**Figure 1A**). Briefly, embryos generated by *in vitro* fertilization were first cultured to the blastocyst stage following the method described before [11, 12]. At blastocyst stage (day 6.5), embryos were transferred to culture dishes plated with human primary endometrial cells. At day 8, all cocultured embryos hatched out from zona pellucida, attached to the bottom of the dish and adopted a flattened structure that is very similar to previous reports [11, 12] (**Figure 1B-1G**).

**Figure 1:**
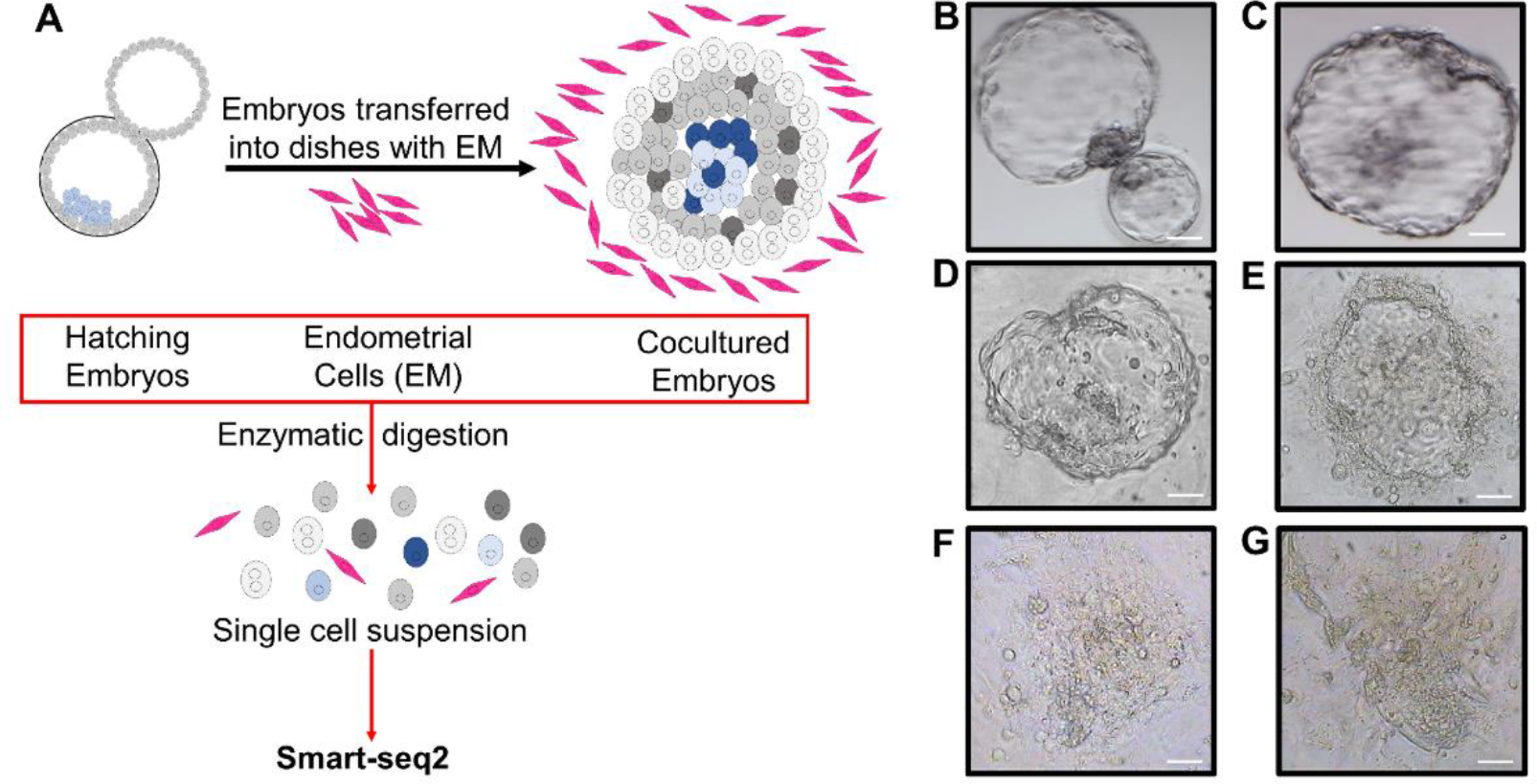
Experiment design of the study. **A:** Human blastocysts generated using *in vitro* fertilization were cocultured with human primary endometrial cells. Endometrial cells and embryos from blastocysts were dissociated and individual cells were collected for single cell RNA sequencing using SMART-seq2 protocol. **B-G**: Representative microscopic images of human peri-implantation embryos, including day 6 embryos **(B),** day 7 un-cocultured **(C)** and cocultured **(D)** embryos, day 8 embryos **(E),** day 9 embryos **(F)** and day 10 embryos **(G).** (Scale bars = 100μm)

To obtain transcriptomic profiles of human trophoblast cells during peri-implantation development, we harvested single cells from 19 embryos from day 6 to day 10, complement with 25 endometrial cells. Transcriptomes from 614 single cells were successfully profiled, with 0.7 million uniquely mapped reads and 24,011 detected transcripts per cell on average. Principle component analysis (PCA) and unbiased hieratical clustering showed that embryonic cells and endometrial cells form two distinct clusters (**Figure 2A**), and some endometrial cells were mislabeled as embryonic cells during cell picking (**Figure 2B**). Therefore, we excluded all endometrial cells and focused on 516 embryonic cells in our subsequent analysis.

**Figure 2:**
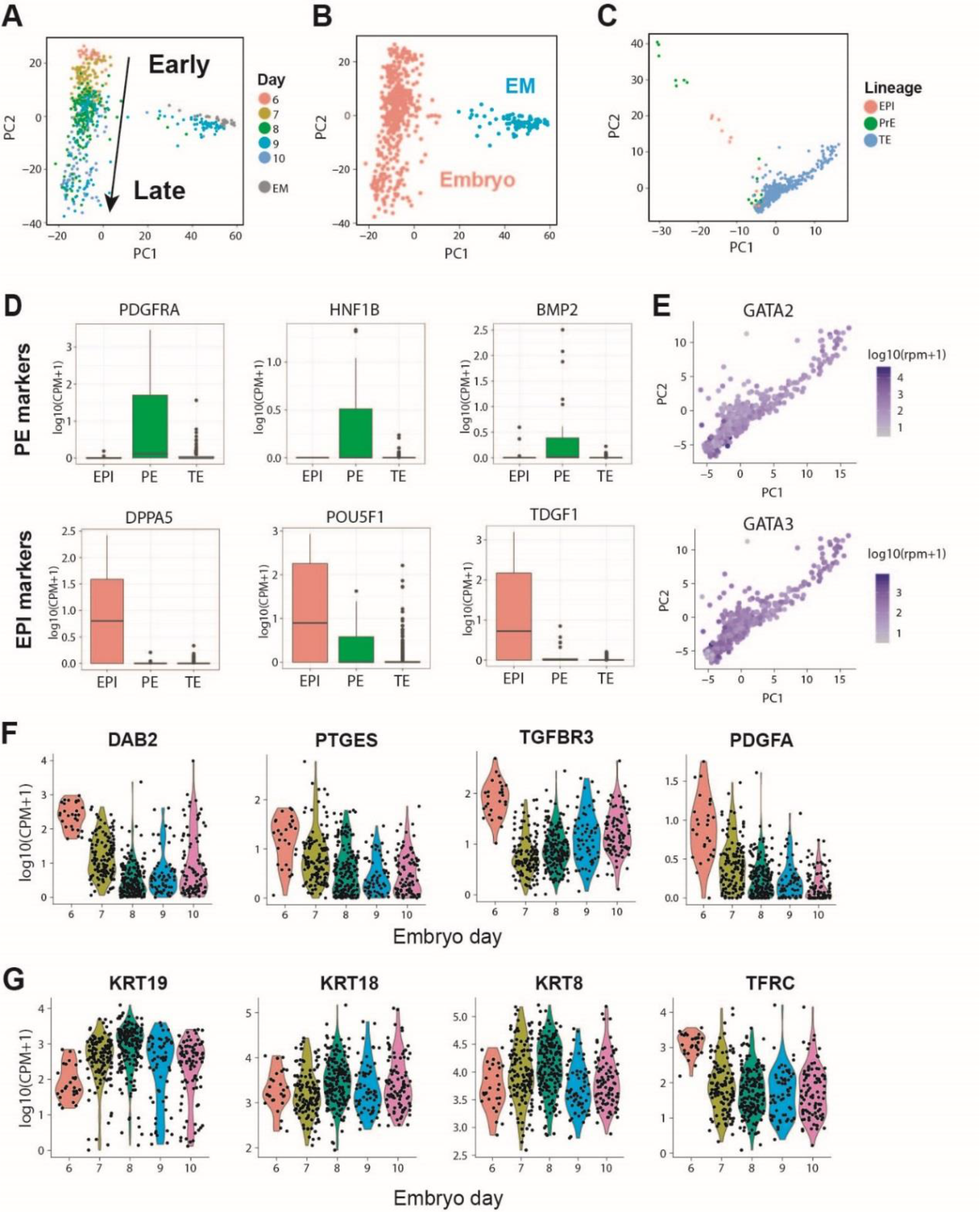
Single-cell RNA sequencing revealed maintained marker genes for trophoblasts during peri-implantation development. **A:** Principle component analysis showed that cells form two distinct clusters, representing embryonic cells and endometrial cells. **B:** Some endometrial cells were annotated as embryonic cells during manual cell picking. Hierarchical clustering using transcriptome data unbiasedly separated embryonic cells and endometrial cells, and the new cell identities were visualized on PCA. **C:** Embryonic cells were classified into EPI-, PE- or TE-lineage cells based on their expression of 300 previously defined lineage markers. The cell lineage identities were visualized on PCA computed using the 300 lineage marker genes. **D**: Boxplot showing the expression of representative lineage markers in EPI- or PE-lineage cells we defined. **E:** Scatter plot showing the two classic TE marker genes, namely *GATA2* and *GATA3*, were sustainably expressed in TE and trophoblasts during peri-implantation development. **F:** Violin plot showing the expression of 4 previously reported TE-lineage markers during peri-implantation development. **G:** Violin plot showing the expression of 4 highly expressed TE-lineage markers we identified during peri-implantation development.

The 516 embryonic cells consist of cells of epiblast (EPI), primitive endoderm (PE) and trophectoderm (TE) lineages. To separate trophoblasts that derived from TE lineage from EPI and PE lineage cells, we determined lineage origination of each cell based on its expression of 300 lineage markers, using an algorithm reported before [13]. We identified 476 TE-, 14 EPI- and 26 PE-lineage cells. PCA analysis using the 300 lineage markers showed that EPI and PE cells separated from TE-lineage cells (**Figure 2C**). In addition, EPI and PE cells highly express their corresponding maker genes (**Figure 2D**), whereas well-characterized TE markers such as *GATA2* and *GATA3* were highly expressed in all TE-lineage cells (**Figure 2E**). These results suggested that this algorithm faithfully separated EPI and PE lineage cells from TE-lineage cells.

Interestingly, while *GATA2* and *GATA3* were highly expressed through day 6 to day 10, the expression of other TE markers such as *DAB2, PTGES*, *TGFBR3* and *PDGFA* were significantly downregulated after implantation at day 7 (**Figure 2F Figure S1A**). These results indicated that not all TE lineage markers are sustainably expressed in trophoblasts during peri-implantation development. We found additional four TE markers, namely *KRT19*, *KRT18*, *KRT8* and *TFRC*, that were consistently highly expressed in TE and trophoblasts from day6 through day 10 using our data (**Figure 2G, Figure S1B**). Collectively, our single-cell RNA-seq data provided a comprehensive transcriptomic profiling for trophoblast from blastocyst through early-post-implantation stages.

### Weighted Gene Co-expression Network Analysis (WGCNA) Reveals Genetic Program Dynamics for Peri-Implantation Trophoblasts Development

PCA showed that trophoblast cells were grouped by their development day (**Figure 2A**). To systematically investigate the genetic program dynamics, we performed Weighted Gene Co-expression Network Analysis (WGCNA) on 2,464 genes that were variably expressed in trophoblast cells between different developmental stages. WGCNA identified eight gene modules, each of which contains a set of genes that tend to be coexpressed at a certain development stage (**Figure 3A**). By relating module expression to development day, we found these eight modules collectively represent three genetic networks that were specifically upregulated at day 6, day 7-8 and day 8-10 (**Figure 3B**). These networks could represent core genetic programs that operate in the early (pre-implantation), middle (during implantation) or late (post-implantation) stage of trophoblasts development. We then performed gene ontology (GO) analysis on genes in each network to investigate their biological function. The early-stage network was enrichment in functional terms related to epithelium-like trophoblast development, including morphogenesis of embryonic epithelium, epithelial cell development and epithelial tube formation. Human endogenous retrovirus (HERV) genes are highly expressed in placenta and have an important function in ST formation [14]. The middle-stage network is significantly associated with RNA catabolic process, viral gene expression and viral transcription, indicating *HERV* genes are dynamically regulated at day 7-8. Genes in the late-stage network were specifically upregulated in day 8-10. GO terms such as cell migration, extracellular matrix organization and response to hypoxia were enriched in these genes, suggesting the activation of trophoblast invasion [15] (**Figure 3C and Figure S2**).

**Figure 3:**
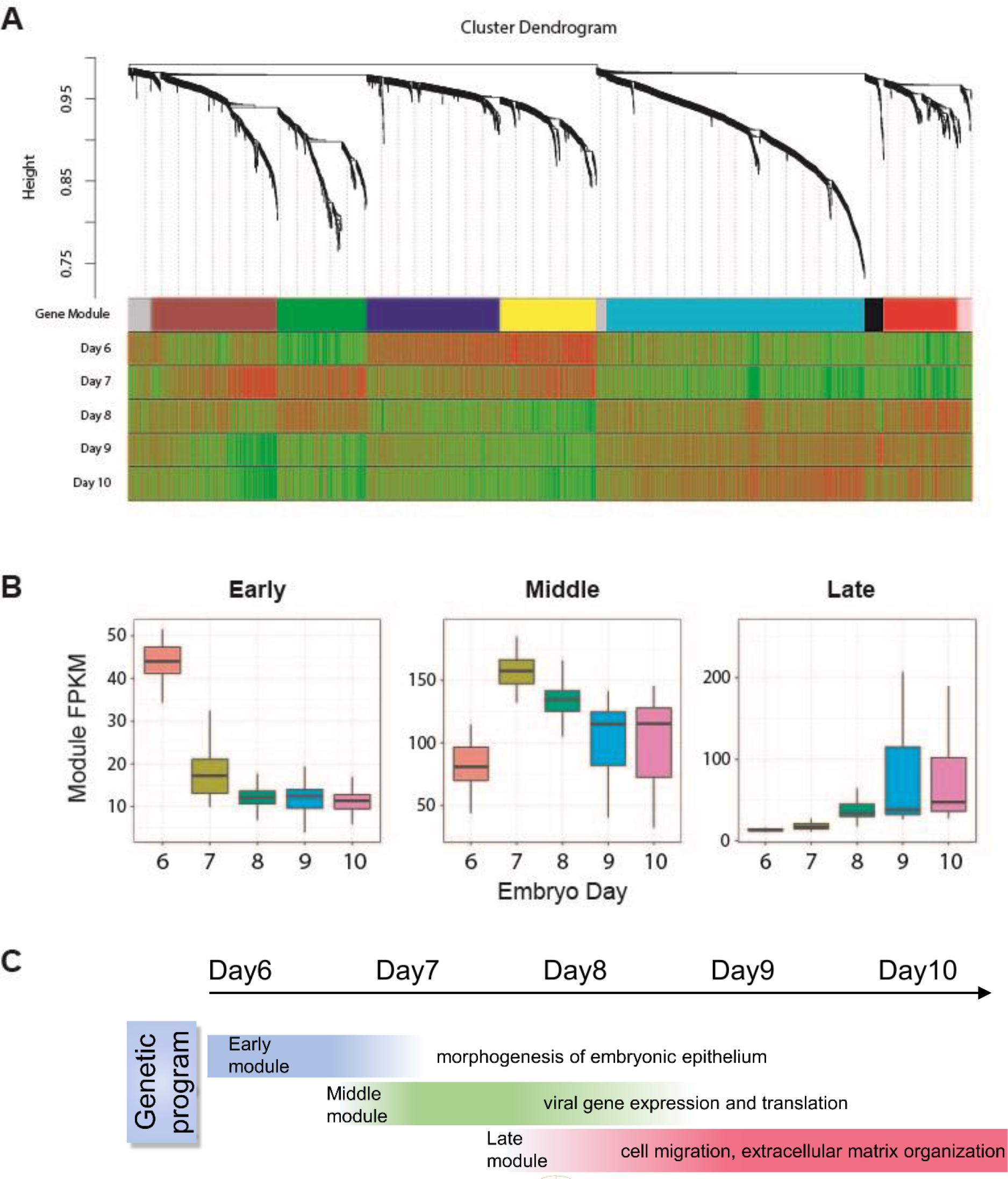
WGCNA revealed genetic networks related to trophoblast development. **A**: Dendrogram showing the gene co-expression network constructed using WGCNA. The color bar labeled as “Gene Module” beneath the dendrogram represents the module assignment of each genes. The remaining color bars represent the correlation of genes with each developmental stages. Red means a gene is positively correlated with a developmental stage, therefore trend to be upregulated at this stage; green means a gene is negatively correlated with a developmental stage therefore trend to be downregulated at this stage. 8 modules were classified into 3 genetic networks that activated at different developmental stages. **B**: Boxplots showing the distribution of network expression (mean RPKM of all genes within a given network) for different developmental stages. **C:** Schematic drawing of the sequential transcriptome switches of representative GO pathways within each network.

To further identify the genes that might play a critical regulatory role in these three genetic networks, we identified 240 hub genes based on the WGCNA measure of intramodular gene connectivity (kME). Hub genes are genes that centrally located within a gene module (kME > 0.8, P<0.05) and have co-expression relationship with many other genes, therefore could have critical regulatory functions. We found that many hub genes are related to critical placental function. For example, *EMP2* was a hub gene in the early-stage network. *Emp2*-deficiency in mice causes aberrant placental angiogenesis [16]. *ESRRG* was identified as a hub for the late-stage network, and abnormal reduction of *ESRRG* expression in human placenta is associated with intrauterine growth restriction and pre-eclampsia [17]. These results suggested many WGCNA hub genes are potential key regulatory genes for early placental development.

### Single-Cell RNA Sequencing Revealed the Timing of Trophoblast Differentiation

Trophoblast sublineages such as EVT, CT and ST are derived from multipotent trophoblasts. But when multipotent trophoblasts differentiate into trophoblast sublineages is unclear [1]. To study this, we performed unbiased clustering using an SNN graph-based clustering algorithm on 476 trophoblast cells, and classified them into six subpopulations (**Figure 4A-B and Figure S3A-S3B**). Cluster 2, 4 and 5 together contain all cells from day 9, day 10, and a few cells from earlier days. By examining the expression of previously defined sublineages marker genes, we found that EVT or ST markers highly expressed in Cluster 2 or 5, whereas Cluster 4 co-express CT and EVT marker genes similar to Cluster 1 (**Figure 4C**). We then identified genes that were specifically expressed in Cluster 2, 4 and 5. We found that many ST marker genes, such as *HSD3B1*, *CYP19A1*, *SDC1*, *ERVW*-1 (*Syncytin*-1), *ERVV*-1, *CGA* and *CGB*, were specifically highly expressed in Cluster 5. CT markers, such as *ITGA6* and *FZD5*, were specifically expressed in Cluster 4. A few EVT marker genes such as *MMP2* were specifically highly expressed in Cluster 2 (**Figure 4D**). Taken together, these results suggest that Cluster 5, Cluster 4 and Cluster 2 represent ST, CT and EVT respectively. The rest clusters consist of trophoblasts from day 6 through day 8, and express CT markers. These results indicated that these clusters represent multipotent trophoblasts that have not committed to differentiation.

**Figure 4:**
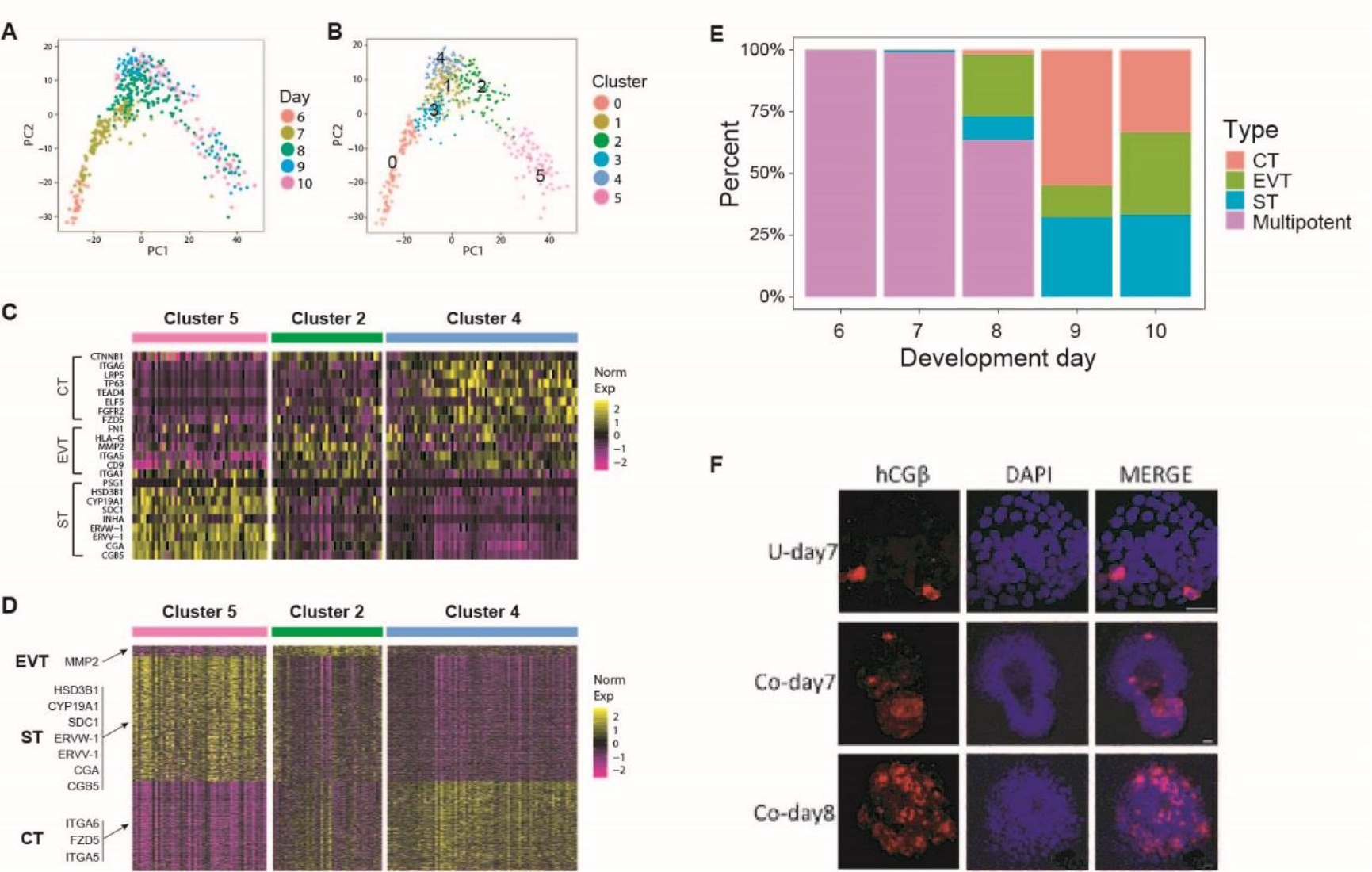
Single cell RNA-seq revealed the timing of trophoblast differentiation. **A-B**: Unbiased clustering based on SNN graph classified trophoblasts into 6 subpopulations. Cells at day 9 and day 10 were classified into 3 subpopulations. **C**: Heatmap showing the expression of previous defined ST, EVT or CT marker genes within the 3 subpopulations identified from day 9 and day 10 cells. **D**: Heatmap showing the expression of genes that were specifically expressed in within the 3 subpopulations identified from day 9 and day 10 cells. We therefore annotated these 3 subpopulations as CT, EVT and ST, respectively. **E**: Stacked barplot showing the percentage of the CT, EVT and ST at different development day. The ST first appear at day 7 in cocultured embryos, while the EVT first appear on day 8. **F**: Immunostaining of ST marker gene hCGβ in embryos at different stages. (Scale bars = 100μm)

We then tried to determine when EVT and ST were established in embryos. We found that ST first appears in cocultured embryos at day 7, even though at a very low percentage (1 out of 60 cells). ST cells then become more abundant at day 8 (16 out of 168 cells) (**Figure 4E**). Immunostaining showed that hCGβ positive cells can be detected as early as day 7, and become more abundant at day 8 (**Figure 4F**). These results suggested that ST cells first occur after day 7, and become more abundant after day 8. Similarly, we found EVT cells were absent in all of the day 7 embryos but appear at day 8, indicating EVT were generated after day 7 (**Figure 4E**). Taken together, our results revealed the time course when ST and EVT appear in peri-implantation embryos *in vitro*.

### Single-Cell Bifurcation Analysis Using the Variance of Gene Expression (SCBAV) Identifies TBX3 as a Novel Master Regulator for Trophoblast Differentiation

The ST is an important trophoblast sublineage that forms the primary barrier between maternal and fetal circulation and synthesize hormones vital for pregnancy. The ST is derived from multipotent trophoblasts within TE and CT. Previous studies using mature placentas and cell lines have demonstrated that many regulatory factors and pathways have been reported to be linked with the human ST formation [18-21]. However, these results must be interpret carefully, because these *in vitro* differentiation models may not perfectly recapuliate the mechanism for trophoblast differentiation *in vivo*.

Our data provided a unique opportunity to study how trophoblast differentiation is regulated *in vivo*, especially during early placenta development. It is generally accepted that the mechanism underlying a cell-fate decision event can be summarized by a hieratical model: a small number of “master” regulators, such as transcriptional factors, were stochastically activated in a subset of cells before cell-fate decision due to environmental fluctuations or random variation of transcriptional network. These master regulators then activate a larger number of cell-type specific genes, which initiate the fate transition and confer cell-type specific function. The identification of the master regulator can be a critical step for understanding the molecular mechanism controlling the trophoblast differentiation. However, the lack of a computational method that can systematically identify master regulators underlying a cell fate decision event poses a major challenge for our analysis.

To identify the master regulator genes that drive the cell fate transition from multipotent trophoblast to ST, we designed single-cell bifurcation analysis using variance of gene expression (SCBAV), a computational strategy that can systemically screen for master regulators using time-serial scRNA-seq data (**Figure S4A**). Briefly, SCBAV first reconstruct a cell development trajectory from scRNA-seq data. A cell type transition will be represented as a bifurcation event in the trajectory. SCBAV then screen the master regulator underlying a bifurcation event according to gene expression level and variation. Transcription factors that are highly variable before bifurcation, and significantly differentially expressed in two lineages after bifurcation are likely to be the master regulator. By applying SCBAV on our trophoblast cells dataset, we identified a bifurcation event that happens after day 8 (**Figure S4B-S4C**). CT and EVT markers were highly expressed in one lineage after bifurcation, while ST markers were greatly upregulated in the other lineage (**Figure S4D-S3I**). This bifurcation, therefore, captured the cell-fate segregation of ST from CT and EVT. SCBAV found 26 putative master regulators driving this bifurcation. Among them, TBX3 is the transcription factor that was ranked as the most top by both of two master regulator screening criteria (**Figure S4J-S4L**). TBX3 is highly expressed in embryos at day 8 (**Figure S5**). Taken together, these results suggested TBX3 could be a master regulator controlling multipotent trophoblast differentiation into ST.

### TBX3 is Required for Differentiation of Trophoblast Cells into Syncytiotrophoblast

To validate the function of TBX3 in trophoblast differentiation, we used a well-established *in vitro* trophoblast differentiation system, which modeled CT-to-ST differentiation by treating trophoblast cell line JEG-3 with cAMP analog 8-Br-cAMP. The generation of ST can be characterized by cell fusion and ST marker gene expression [20, 22]. In control JEG-3 cells that were not treated by 8-Br-cAMP, the cell fusion ratio was less than 1%, and ST marker hCGβ expression was almost undetectable, indicating ST generation before treatment is minimal (**Figure 5A-B, 0mM group**). After 48h treatment with 8-Br-cAMP, JEG-3 cells exhibited greatly enhanced cell fusion and hCGβ compared to the control group, and the fusion ratio is correlated with 8-Br-cAMP concentration (**Figure 5A-B**). Notably, TBX3 was not expressed in the control group but was significantly upregulated by 8-Br-cAMP treatment, and the upregulation fold is also correlated with 8-Br-cAMP concentration (**Figure 5A and 5C**).

**Figure 5:**
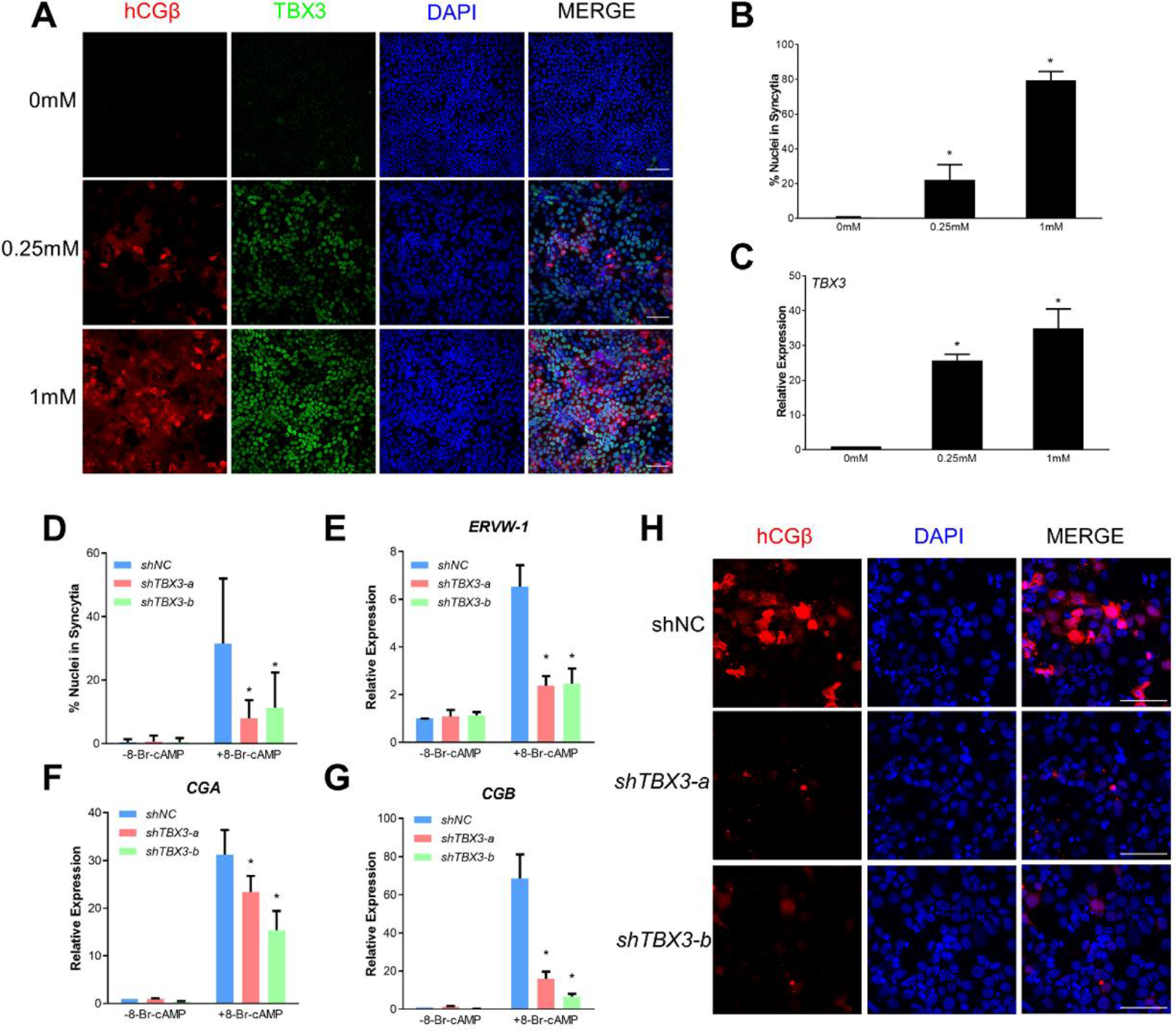
TBX3 regulated trophoblast cell differentiation into ST. **A**: Immunofluorescence images of hCGβ (Red) and TBX3 (Green) showing ST cells formation under 0mM, 0.25mM, and 1mM 8-Br-cAMP for 48h. **B**: Graphical depiction of the percentage of nuclei associated with ST cells following exposure to different concentrations of 8-Br-cAMP for 48h. **C**: qRT-PCR for *TBX3* expression following exposure to different concentrations of 8-Br-cAMP for 48h. **D**: The percentage of nuclei associated with fused cells. **E-G**: qRT-PCR for *ERVW*-1 (**E**), *CGA* (**F**), and *CGB* (**G**) expression in JEG-3 cells expressing *shNC*, *shTBX3-a*, or *shTBX3-b* before and after 0.25mM 8-Br-cAMP treatment for 48h. **H**: Representative images of hCGβ expression in JEG-3 cells expressing *shNC*, *shTBX3-a*, or *shTBX3-b* cultured under 0.25mM 8-Br-cAMP for 48h. *p<0.05, n≥3, Mean ± SD. (Scale bars = 100μm)

We then knocked down TBX3 in JEG-3 cells using lentiviral vectors expressing TBX3 shRNA and assessed its influence on ST generation. Control shRNA (*shNC*) targeting no known mammalian RNA and shRNA targeting TBX3 (*shTBX3*-a and *shTBX3*-b) had no discernable influence on proliferation of JEG-3 cells. However, *shTBX3*-a and *shTBX3*-b interference resulted in 96±1% and 97±1% suppression of *TBX3* mRNA levels after 8-Br-cAMP treatment, respectively and also in suppressed TBX3 protein levels (**Figure S6A-B**). TBX3 knockdown also significantly reduced cell fusion (75±7% and 64±14% reduction in *shTBX3*-a and *shTBX3*-b cells, respectively) (**Figure 5D**) and downregulated ST markers transcription, including human chorionic gonadotropin subunits (*CGA* and *CGB*), *Syncytin* (*ERVW*-1) and other HERV-derived genes (*ERVV*-1 and *ERVV*-2) (**Figure 5E-5G, Figure S6C**). Immunostaining showed that hCGβ protein level was decreased by TBX3 knockdown (**Figure 5H**). Taken together, these results demonstrated that TBX3 is required for ST formation.

**Figure 6:**
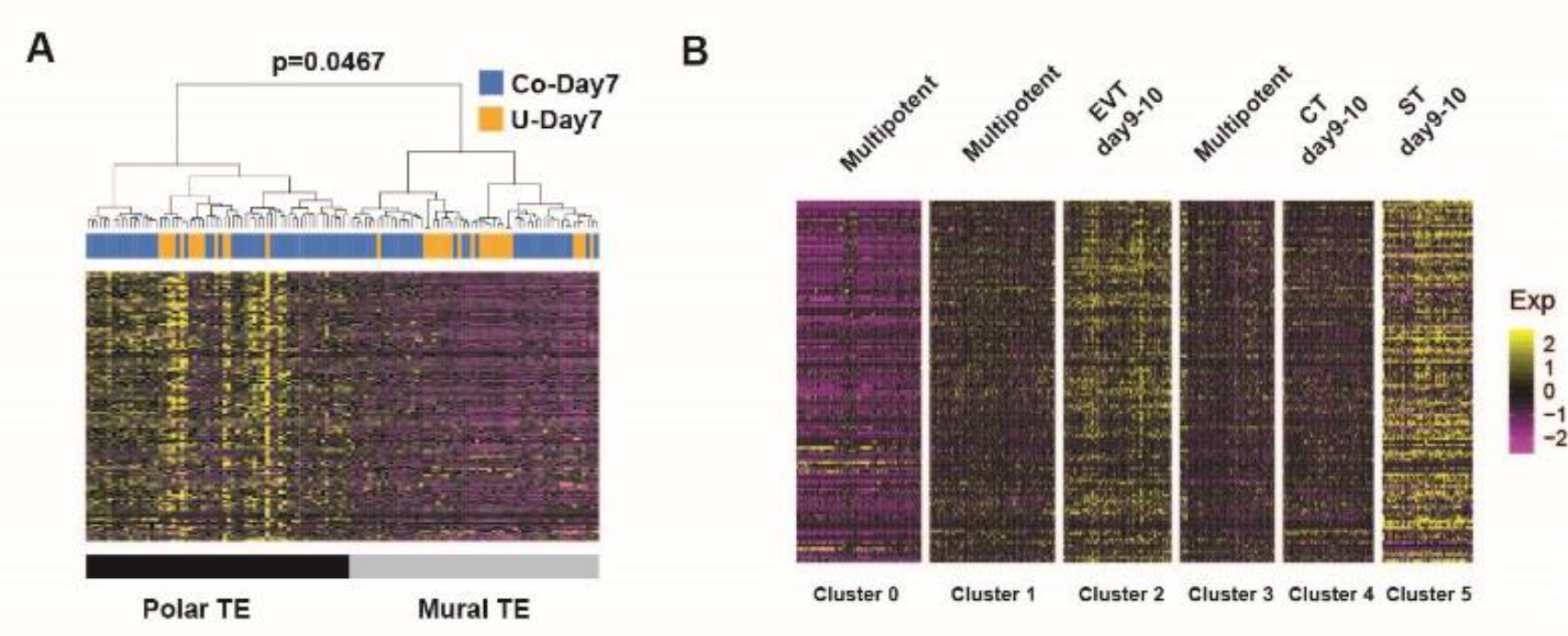
**A**: Hierarchical clustering using previously identified polar-TE markers classified day 7 trophoblasts into 2 clusters with high or low polar-TE markers expression respectively. **B**: Heatmap showing the expression of previously identified polar TE markers at 6 trophoblast clusters identified before.

### Coculturing with endometrial cells influences genes related to trophoblast development

The unbiased SNN-graph clustering assigned trophoblasts from day 7 embryos cocultured with endometrial cells (co-day 7 trophoblasts) and trophoblasts from day 7 embryos cultured without endometrial cells (u-day 7 trophoblasts) into two distinct clusters, indicating that endometrial cells have a profound impact on trophoblasts (**Figure S7**). To dissect the influence of coculturing, we identified the differentially expressed genes (DEGs) between co-day 7 and u-day 7 trophoblasts. We found *EIF5A*, a gene involved in trophoblast proliferation, migration, and invasion; *WEE1*, a gene regulating mitosis and associated with cell cycle progression in trophoblast cells; and *CCR7*, a chemokine gene associated with trophoblast differentiation, were significantly upregulated by coculturing in day 7 trophoblasts [23-25]. On the other side, *MDM2*, a gene associated with preeclampsia susceptibility, was significantly downregulated by coculturing [26]. These results suggest that coculturing with endometrial cells can influence the expression of genes related to trophoblast development.

### Transcriptomic Analysis Suggests Trophoblast Differentiation Initiates at Polar TE Side in Blastocysts

Polar and mural TE cells are TE subpopulations that emerge at the blastocyst stage [13, 27]. Anatomical studies showed that most of human blastocysts attach to endometrium at the polar side, and polar TE first proliferate and invade into endometrium [1, 3]. These results implicated the polar TE could have an important function for implantation. But the role of polar TE during post-implantation development is still elusive. Hieratical clustering using 129 previously identified polar TE markers [13] robustly separate day 7 trophoblast cells into 2 subpopulations, which is consistent with the existence of polar and mural TE at day 7 (**Figure 6A**). We then sought to investigate the relationship between polar and mural TE and the 6 subpopulations we identified above. We found that polar TE markers were lowly expressed in all day 6 cells and day 7 non-cocultured cells, and moderately expressed in CT cells at different developmental day. Interestingly, polar TE markers were significantly upregulated in differentiated trophoblasts, including EVT and ST (**Figure 6B**). These results showed that polar TE markers were upregulated in differentiated trophoblast cells, suggesting that trophoblast differentiation was first initiated in polar TEs.

## Discussion

Trophoblasts undergo magnificent morphological movement and cellular changes after implantation. In this study, by profiling over 500 single cells in 19 embryos generated using a coculture system, we reconstruct the transcriptome dynamics of trophoblasts through blastocyst to early post-implantation stages (**Figure 7**). Our study complements previous studies that use scRNA-seq to profile trophoblasts and other cell types within mature placenta [4-7].

**Figure 7:**
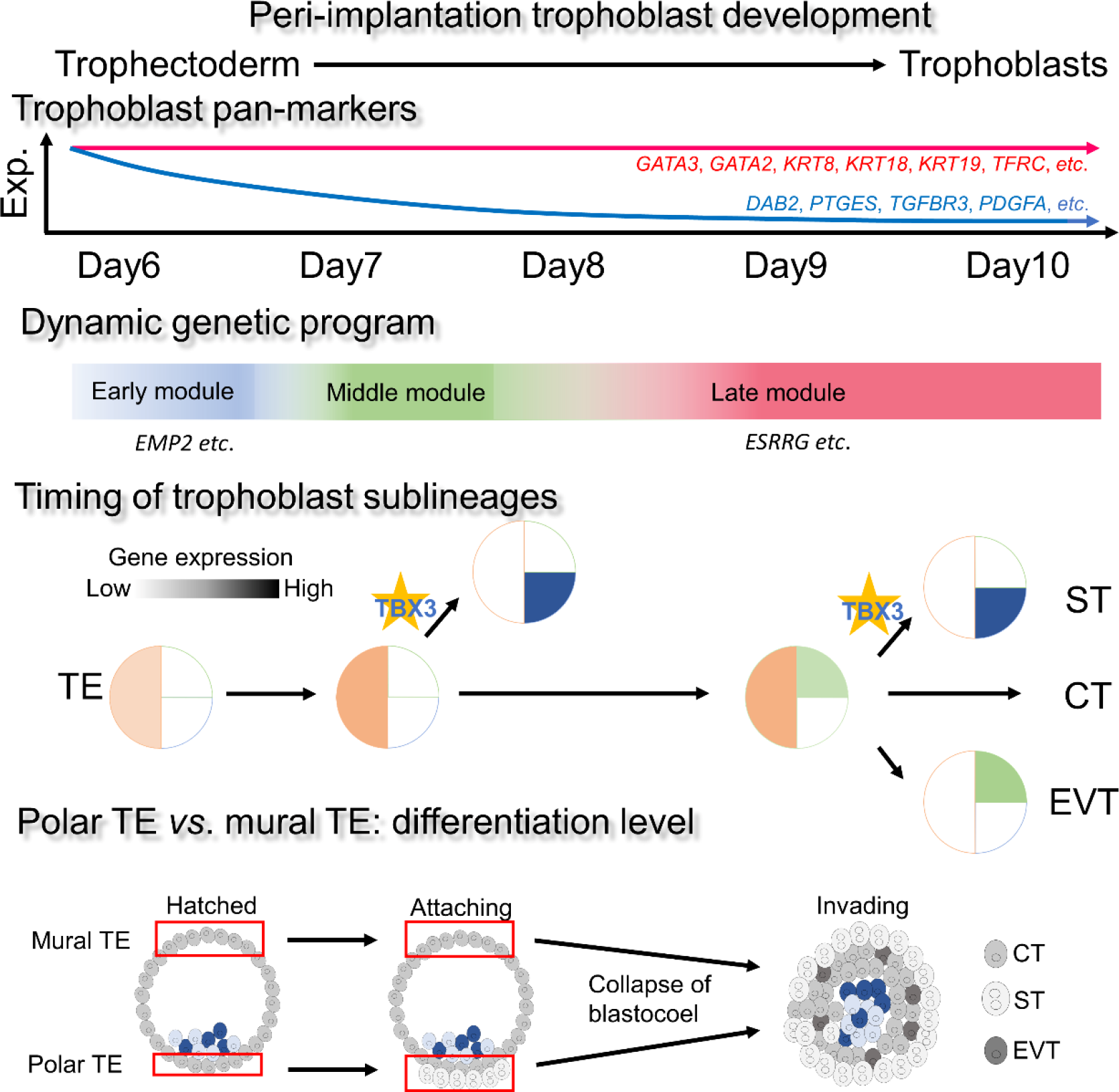
Schematic diagram of trophoblast development during peri-implantation. Our single-cell profiling of peri-implantation trophoblasts revealed the differential expression of trophoblast pan-markers and the dynamic genetic program of trophoblast development. ST cells were abundant at day8, and TBX3 was critical for ST formation using SCBAV as well as *in vitro* verification. Trophoblast differentiation was also found earlier in polar TE cells than mural TE cells.

Our bioinformatic analysis suggests TBX3 has a critical role in regulating the ST formation. TBX3 haploinsufficiency in human causes ulnar-mammary syndrome (UMS), a genetic disorder characterized by abnormal forelimb and apocrine gland development [28]. Although *TBX3* is abundant in human placenta [29], its role in the placenta has not been studied. Our results demonstrated that TBX3 is required for both hCGβ and syncytium formation, indicating TBX3 is a master regulator for ST formation.

HERV-derived genes are upregulated in ST, and their expression is regulated by DNA methylation [30]. Using the in vitro trophoblast differentiation system, we found that *DNMT1* and *DNMT3B* are significantly downregulated after 8-Br-cAMP treatment, whereas *TET2* and *TET3* are significantly upregulated after 8-Br-cAMP treatment (**Figure S6C**). These results are consistent with our scRNA-seq data, and indicate that *DNMT1*, *DNMT3B TET2* and *TET3* might be involved in the regulation of DNA methylation on HERVs during trophoblast differentiation. However, TBX3 knockdown downregulated *DNMT1* but did not significantly influence expression level of *DNMT3B*, *TET2* nor *TET3* (**Figure S6C**). These results indicate that TBX3 can regulate HERV expression but not through altering the transcription level of *DNMT*s and *TET*s. Further studies should be done to elucidate how TBX3 regulates HERV expression in trophoblasts.

Our scRNA-seq data revealed that *DPPA3* (a.k.a *STELLA* and *PGC7*) expression is significantly downregulated in ST compared to CT and EVT (**Figure S8**). Consistently, in the trophoblast *in vitro* differentiation system, *DPPA3* is significantly downregulated by 8-Br-cAMP treatment, whereas TBX3 knockdown does not significantly alter *DPPA3* expression (**Figure S6C**). Since Dppa3 is a factor that safeguards DNA methylation at ERVs in mouse embryos and involves pluripotency circuit in mESC [31], these results suggest DPPA3 in human could maintain differentiation potential of trophoblast progenitors and repress trophoblast differentiation and deactivating HERVs.

In conclusion, our study established single-cell transcriptomic profiles of peri-implantation trophoblast cells. Moreover, we have found a new role of TBX3 as a regulator of the trophoblast differentiation in human. Our study offers unique resources for understanding the early placenta development and pathogenesis associated with early trophoblast defects.

## Materials and methods

### Human Embryo Culture and Isolation

The embryos were voluntarily donated by the patients at the Center for Clinical Reproductive Medicine at Tongji Hospital of Tongji University with informed consent and institutional approval. Human peri-implantation embryos were cultured according to protocols [11, 12] with the addition of endometrium cells. When embryos hatch out from zona pellucida at day 6.5, we transferred blastocysts to dishes plated with human endometrial cells. Non-coculture day7 embryos were used as a control group to coculture day 7. Single cells of embryos randomly collected from all stages were acquired according to a previous procedure [32]. For attached embryos from day 7 to day 10, they underwent short incubation in trypsin to detach from dishes and were manually transferred to a new dish containing trypsin for further dissociation for 3 to 5 minutes with repeated aspiration using a mouth-operated, drawn capillary pipette. A single cell was then manually picked using the capillary pipette into a 0.2-ml PCR tube containing lysis buffer.

### Single-Cell RNA-Seq Library Preparation

Single-cell RNA sequencing was performed using SMART-seq2 protocol with minor modification [33, 34]. Briefly, a single cell was first lysed in 0.5uL lysis buffer, followed by reverse transcription using Superscript III. After purification using Ampure XP Beads, amplified cDNA product was diluted to 0.1ng/uL. 0.1 ng amplified cDNA was used for library construction. Libraries were pooled and sequenced on Illumina Hiseq X10 in paired-end, 150bp mode.

### Single-Cell RNA-Seq Data Processing

Single-cell RNA seq data were first trimmed with TrimGalore! using following parameters “-q 20 --phred33 --gzip --length 30 --paired” to remove adaptor sequences and low-quality bases. The trimmed data was then aligned to human reference genome hg38 using STAR v2.6.0 in the pair-end mode with default parameters. The number of reads mapped to each gene was counted using featureCounts v1.6.2, using Gencode hg38 gene annotation. Customized R scripts and published R packages, including Seurat, were used in subsequent analysis.

### Identification of TE, EPI and PE Lineage Cells and Identification of Peri-Implantation Trophoblast Markers

To our experience, the majority of cells in peri-implantation embryos are of trophoblast lineage. Therefore, random picking ensures us to unbiasedly sample trophoblast population with a few cells of EPI or PE lineage. Embryonic cells were assigned into three lineages, namely TE, EPI and PE, based on their expression of 300 previous identified lineage marker genes. Specifically, the read-count matrix was first normalized and quality-controlled using Seurat. The lineage identity for each cell was then determined using a previous strategy reported before [35]. Briefly, for each cell, a “TE score”, an “EPI score” and a “PE score” were computed using AddModuleScore function implemented in Seurat package, based on its expression of previously identified markers for each lineage respectively. The cell lineage was then defined as the lineage that had the highest score. To identify maintained trophoblast markers, we started with previously identified TE marker genes for pre-implantation embryos, and excluded genes that were lowly expressed in trophoblasts at any stage between day-6 to day-10 (mean FPKM < 10 over all trophoblasts on each day).

### Weighted Gene Co-Expression Analysis (WGCNA)

WGCNA was performed on normalized gene expression data measured in read count per million (RPM) metric, using 2,464 highly variably expressed genes determined by FindVariableGenes function in Seurat. The WGCNA was then performed following the previously published study [32]. Briefly, the topological overlap matrix (TOM) was constructed with softPower was set to 8. The hub genes for each module were identified as module eigengene based connectivity kME > 0.8 and P-value <0.05. The gene ontology enrichment analysis was performed using Gene Ontology Consortium website (http://geneontology.org) and R package clusterProfiler.

### Single-Cell Bifurcation Analysis Using Variance of Gene Expression (SCBAV)

SCBAV consists of two steps. It first constructs a cell lineage trajectory using scRNA-seq data and identifies when lineage segregation happens. It then identifies the master regulator that could cause the lineage segregation based on certain criteria. Briefly, SCBAV represents all cells from different time points on the same reduced dimension using principle component analysis. The first N principle components are used, such that N is the smallest number that can capture at least 40% of the total variation within the data. Next, within this reduced dimension SCBAV identifies cell subpopulations within cells from each time-point, using Gap Statistics and K-means clustering. The Gap Statistics is used to determine the number of subpopulations within cells from each time-point, and K-means clustering is used to classify cells into clusters based on the cluster number determined by Gap Statistics. SCBAV then constructs cell lineage trajectory by iteratively connecting the most similar clusters between two time points. Specifically, for a cluster A in time-point t, SCBAV connects it to the cluster in t+1, whose centroid as the highest correlation with cluster A’s centroid among all clusters in t+1. SCBAV keeps doing this until all clusters from all time-points were connected to one lineage trajectory.

A cell-fate segregation event will be represented as a bifurcation on the cell-fate trajectory. Suppose there is a bifurcation event in the lineage trajectory right after time-point t. Bifurcation event represents a cellular differentiation event. SCBAV then tries to identify the master regulator than drive the bifurcation. According to the hieratical model for cell-fate transition regulation mentioned above, a master regulator that promotes a cell lineage bifurcation must meet two criteria: 1) it is consistently differentially expressed between two lineages after bifurcation point; 2) it is highly variably expressed in the cell population right before bifurcation.

### Cell Culture and shRNA Constructs

Endometrium tissue was obtained from patients at Tongji Hospital of Tongji University. The endometrium tissue donor with no genetic disorder or diseases signed informed consents. Primary endometrium was dissected using scalpels and then enzymatically digested in 1% collagenase I and IV (Gibco,17100-017 and 17014-019) with repeated pipetted up and down for 30 minutes at 37°C. After filtration through 40μm strainers and centrifugation at 300 g for 3 min, the supernatant was removed, and the pellet at the bottom was washed with PBS twice. Then, the pellet was resuspended and prepared for use.

JEG-3 trophoblast cells were purchased from Cell Bank of Chinese Academy of Sciences and were cultured in MEM medium (Gibco, 10370021) with the addition of 0.11g/L Sodium Pyruvate (Gibco, 11360070) and 10% FBS (Gibco, 10099141). To stimulate JEG-3 trophoblast differentiation, cells were treated with 8-Br-cAMP (Sigma-Aldrich, B7880).

Human *TBX3* shRNA and control shRNA constructs in pLenR-GPH vectors were purchased from Shanghai Taitool Bioscience Co., Ltd. (China). The target sites for the shRNAs are *TBX3*-a, GCGAATGTTTCCTCCATTTAA; *TBX3*-b, GCAGTCCATGAGGGTGTTTGA; Control, TTCTCCGAACGTGTCACGT. Lentiviruses were also generated in Shanghai Taitool Bioscience Co., Ltd. (China). For *TBX3* knockdown experiment, lentivirus transduction at a multiplicity of infection (MOI) of 10 was accomplished according to previous studies [36, 37]. Transfected Cells were treated with the addition of 2μg/mL puromycin (Gibco, A1113803) for 2-3 passages and cells stably expressing GFP were used for subsequent analyses.

### Immunofluorescence Staining

Embryos and cells growing on coverslips were fixed in 4% paraformaldehyde and were permeabilized in 0.2% triton X-100 for 10 min. Subsequently, embryos and cells were blocked at room temperature in 5% BSA in PBS for 1 hour and incubated with primary antibodies (hCGβ, Abcam, ab9582; TBX3, Abcam, ab99302) in blocking solution overnight at 4°C. The embryos and cells were then washed twice in blocking solution and incubated with species-appropriate fluorescent-conjugated secondary antibodies at RT for 1 h before final washes in blocking solution. Coverslips with embryos or cells were then moved to drops of Vectashield mounting media with DAPI (Vector Lab, H-1200) on slides for 10 min incubation before imaging. For non-cocultured embryos, all immunostaining operations were under the stereoscopic microscope. Cell fusion index was analyzed according to previous reports [20, 22].

### qRT-PCR

Total RNA was extracted from cultured cells using Takara MiniBest Universal RNA Extraction Kit (Takara, 9767). cDNA was synthesized through RevertAid First Strand cDNA Synthesis Kit (Thermo Scientific, K1622). qRT-PCR was carried out using TB Green™ Premix Ex Taq™ II (Takara, RR820A) and Roche Light Cycler 96 system (Roche). Relative levels of RNAs were calculated by the ddCt method with *GAPDH* as endogenous controls. The primers were shown in Table S1.

### Ethics Statement

All the embryos and endometrium tissue donors signed informed consent. All the procedures were approved by the Institutional Review Board (IRB) of Tongji Hospital in Tongji University.

### Availability of data and material

All sequencing data generated in this study are available on Gene Expression Omnibus (GEO) with accession number GSE125616 (Reviewer access token: ydmncaggxbqdhop). All intermediate data, results, and code used in the analysis are available upon reasonable request to the authors.

## Author contributions

B.L., Q.A., G.F. and Z.X. designed the study. Q.Z., B.L., Y.J. and Z.X. assisted with patient consent, embryos culture and sample dissociations. B.L., P.L., Q.Z., X.Z., and Y.J. performed wet-lab experiments. Q.A. performed the computational analysis.

B.L. and Q.A. interpreted the data and drafted the manuscript. All authors contributed to the manuscript.

## Acknowledgement

This work was supported by grants from the National Natural Science Foundation of China (81771651, 81873832, Key Program 81430026) and National Key R&D Program of China (2017YFC1001301, 2016YFC1000208). Additional support was from Basic Research Projects of Shanghai Science and Technology Commission (16JC1404700).

## Supplementary materials

**Figure S1:**
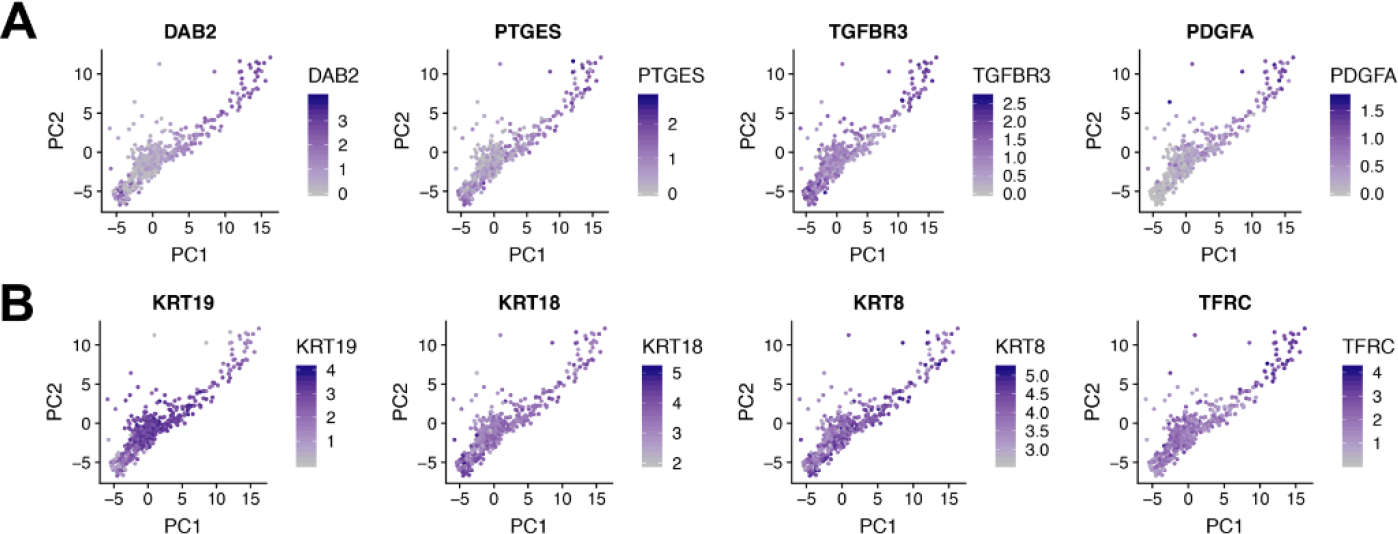
Scatterplot of TE-lineage markers identified previously and in our study. **A**: Scatterplot showing the expression of previously identified TE-lineage markers. **B**: Scatterplot showing the expression of highly expressed TE-lineage markers identified in this study.

**Figure S2:**
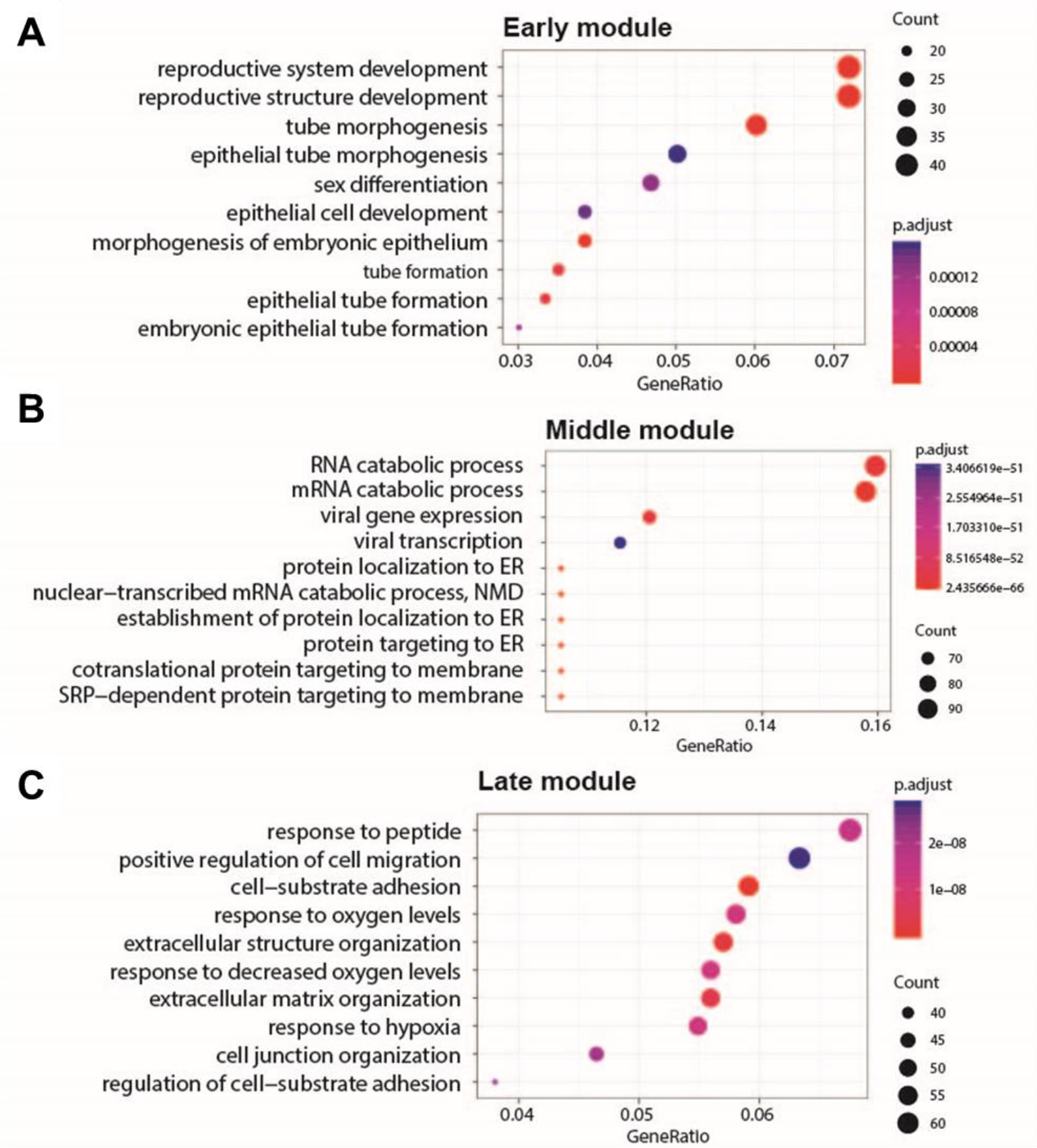
Bobble plot of the gene ontology enrichment of genes within each network.

**Figure S3:**
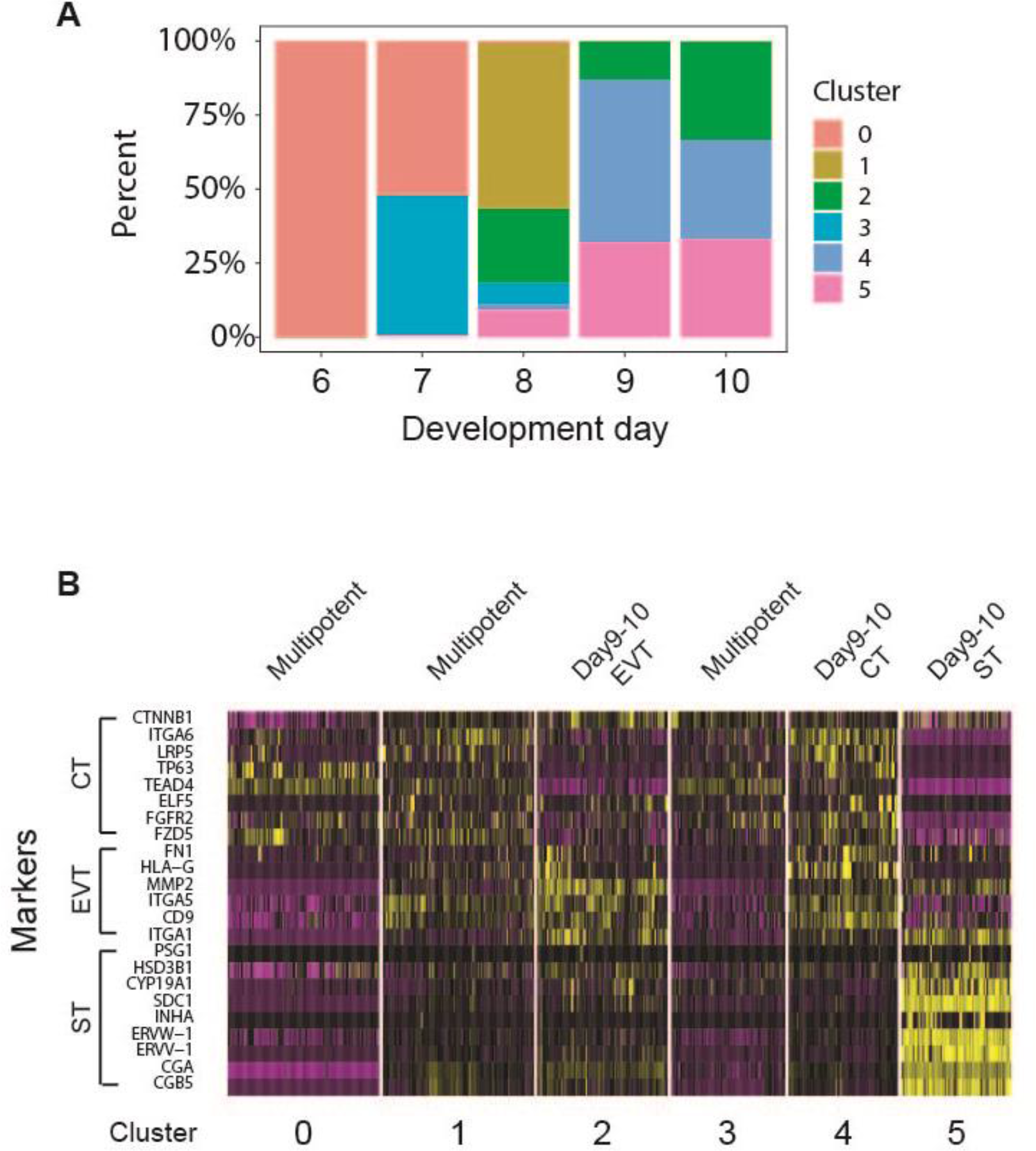
Single-cell RNA-seq revealed the clusters of trophoblast across all development days. **A**: Stacked barplot showing the parentage of cells of 6 subpopulations at different development day. **B**: Heatmap showing the expression of previously identified CT, EVT and ST markers in 6 trophoblast subpopulations.

**Figure S4:**
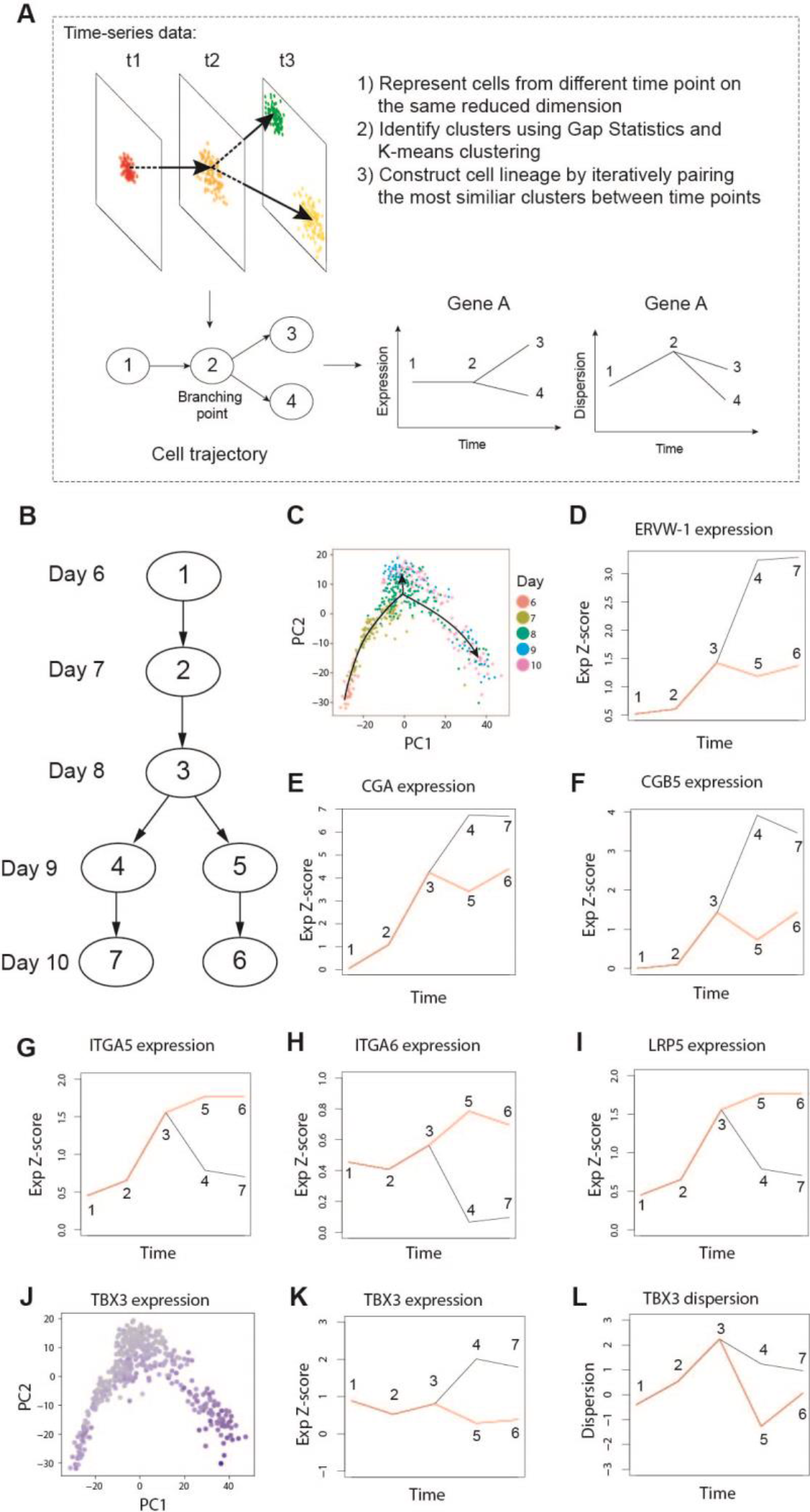
SCBAV Identified TBX3 as a Novel Master Regulator for Trophoblast Differentiation. **A**: Graphical abstract of SCBAV. **B**: Cell trajectory reconstructed by SCBAV. **C**: The bifurcation within the SCBAV cell trajectory recapitulated the cell fate divergence of ST from CT and EVT. **D-F**: Expression of ST specific genes within two lineage branches. **G-I**: Expression of CT specific genes within two lineage branches. **J-L**: TBX3 is variably expressed before bifurcation point, and significantly upregulated in ST compared to EVT and CT after bifurcation.

**Figure S5:**
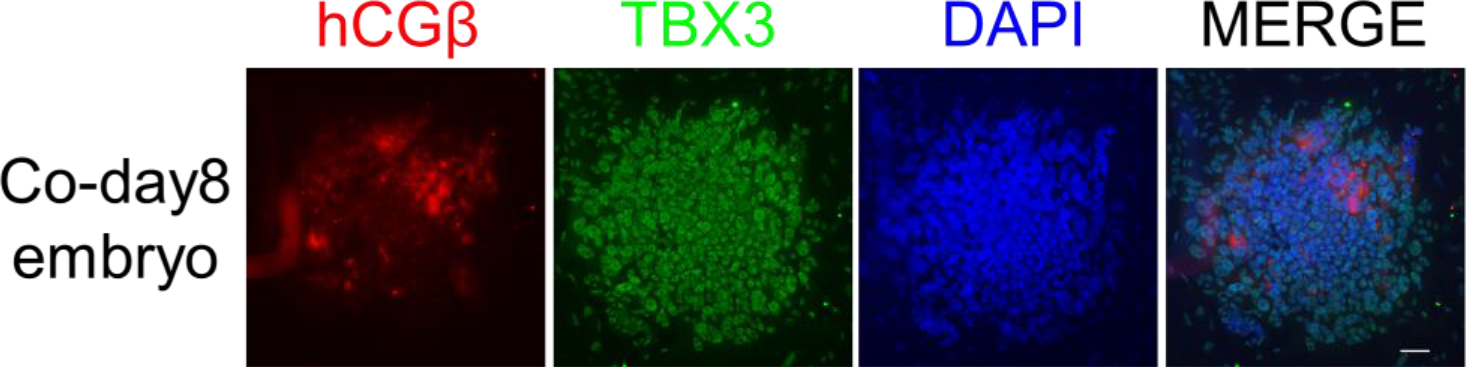
Immunostaining of hCGβ and TBX3 in day8 embryos. (Scale bars = 100μm)

**Figure S6:**
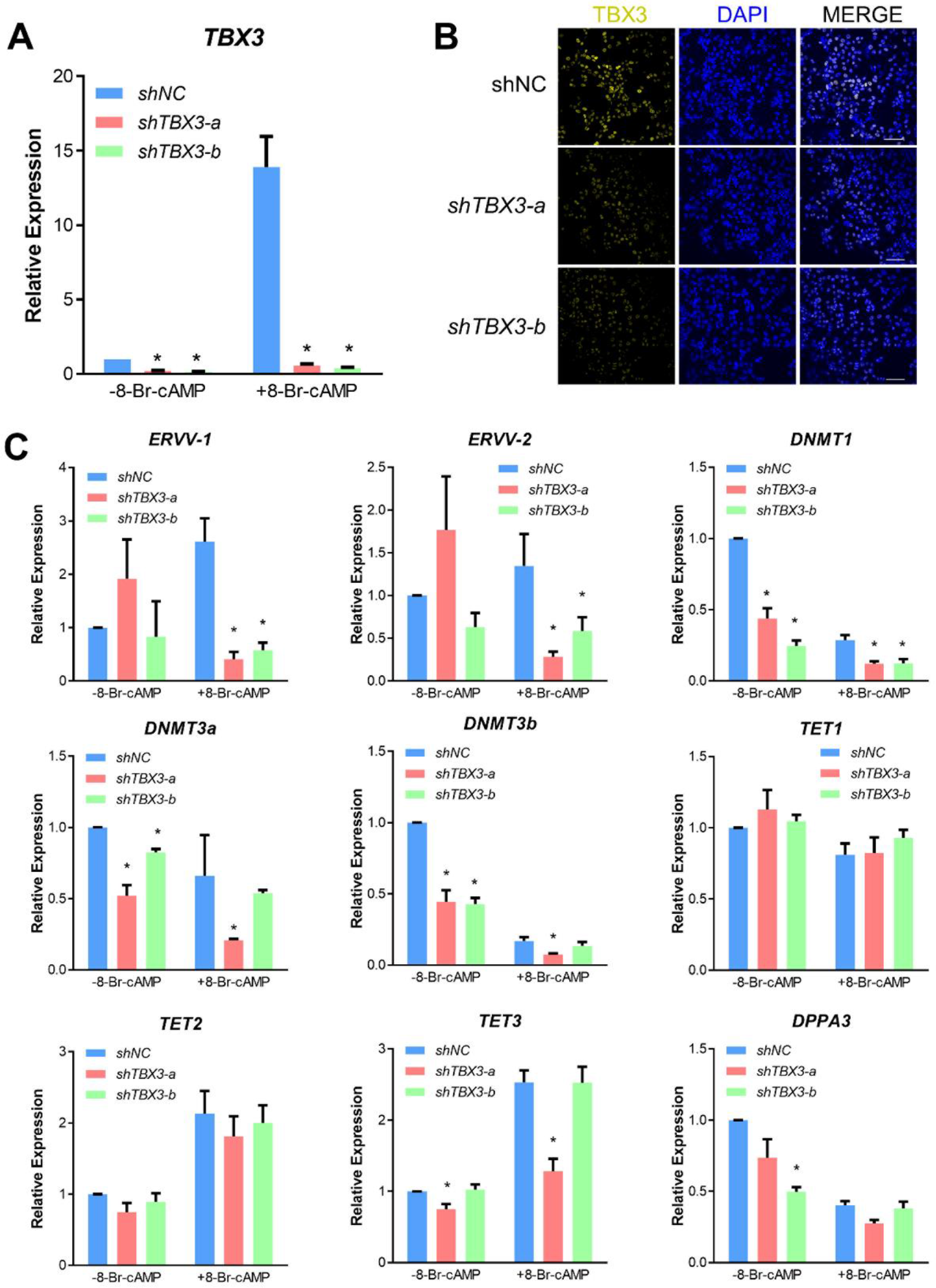
TBX3 regulated trophoblast cell differentiation. **A** and **C**: qPCR for *TBX3*, *ERVV*-1, *ERVV*-2, *DNMT1*, *DNMT3a*, *DNMT3B*, *TET1*, *TET2*, *TET3*, and *DPPA3* expression in JEG-3 cells expressing shNC, *shTBX3*-a, or *shTBX3*-b before or after 0.25mM 8-Br-cAMP treatment for 48h. *p<0.05, n≥3, Mean ± SD. **B**: Representative images of TBX3 expression in JEG-3 cells expressing shNC, *shTBX3*-a, or *shTBX3*-b cultured under 0.25mM 8-Br-cAMP for 48h. (Scale bars = 100μm)

**Figure S7:**
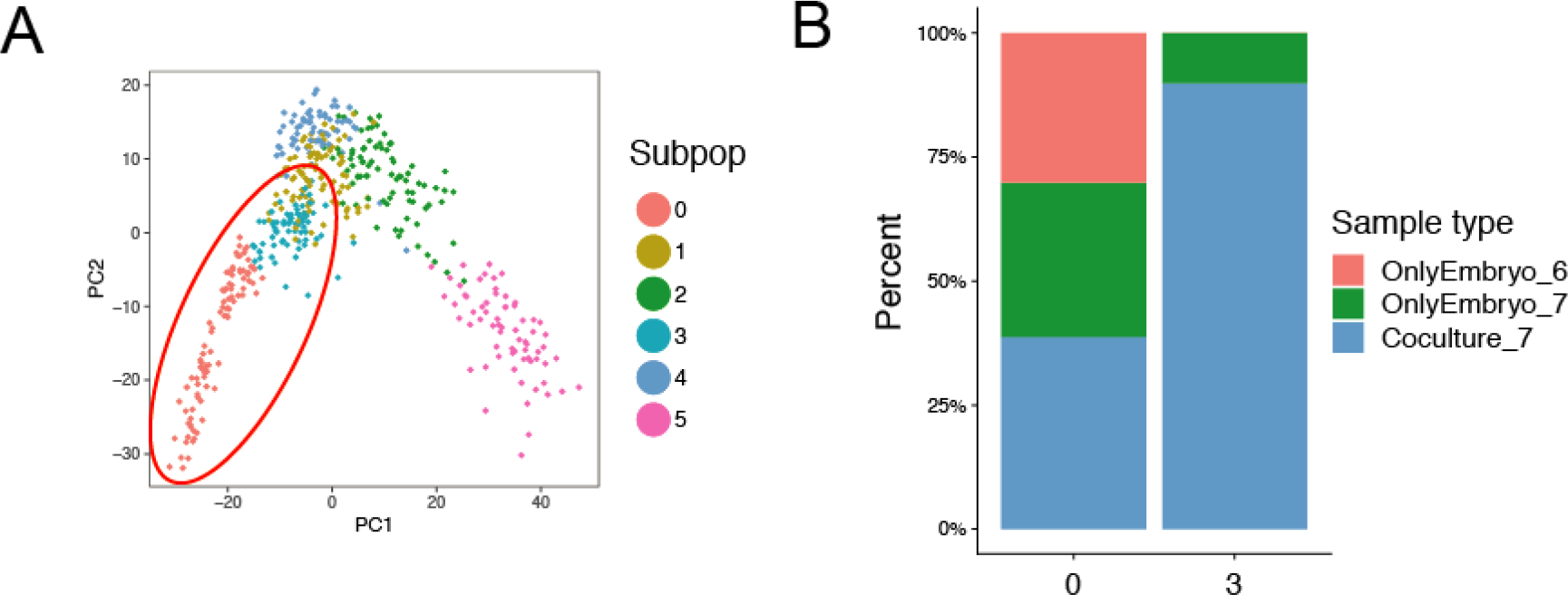
Coculturing induced transcriptomic changes related to trophoblast development. **A**: PCA showing the 6 subpopulation within all trophoblasts. Cells from day 6 and day 7 were assigned into subpopulation 0 and 3. **B**: Stacked barplot showing the percentage of cells of different day and culture condition in subpopulation 0 and 3.

**Figure S8:**
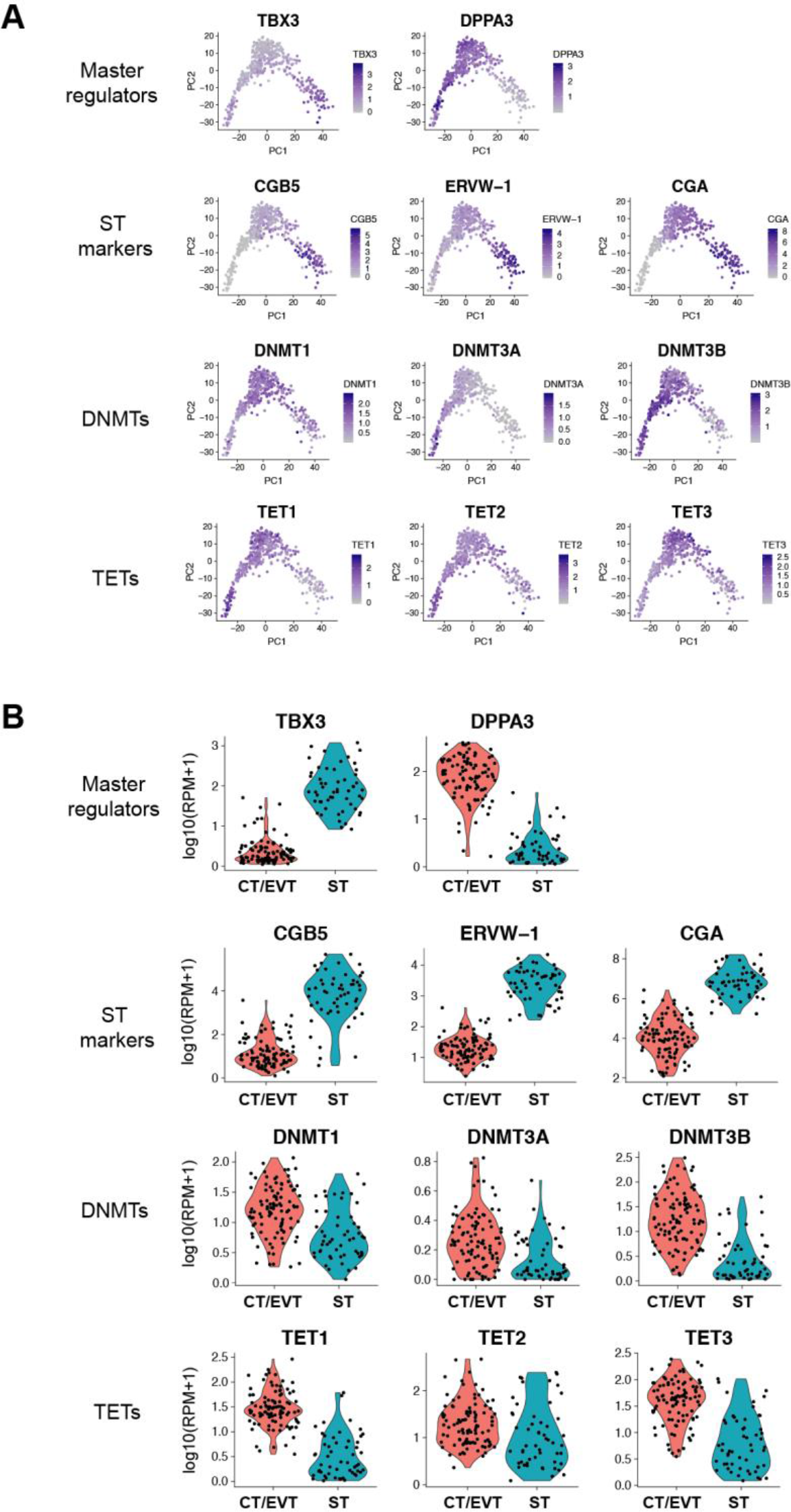
Genes expressed differentially in peri-implantation trophoblast lineages. **A-B**: Scatter plot and violin plot showing the expression of master regulators, ST marker genes DNA methyltransferases and TET methylcytosine Dioxygenases.

**Table S1:**
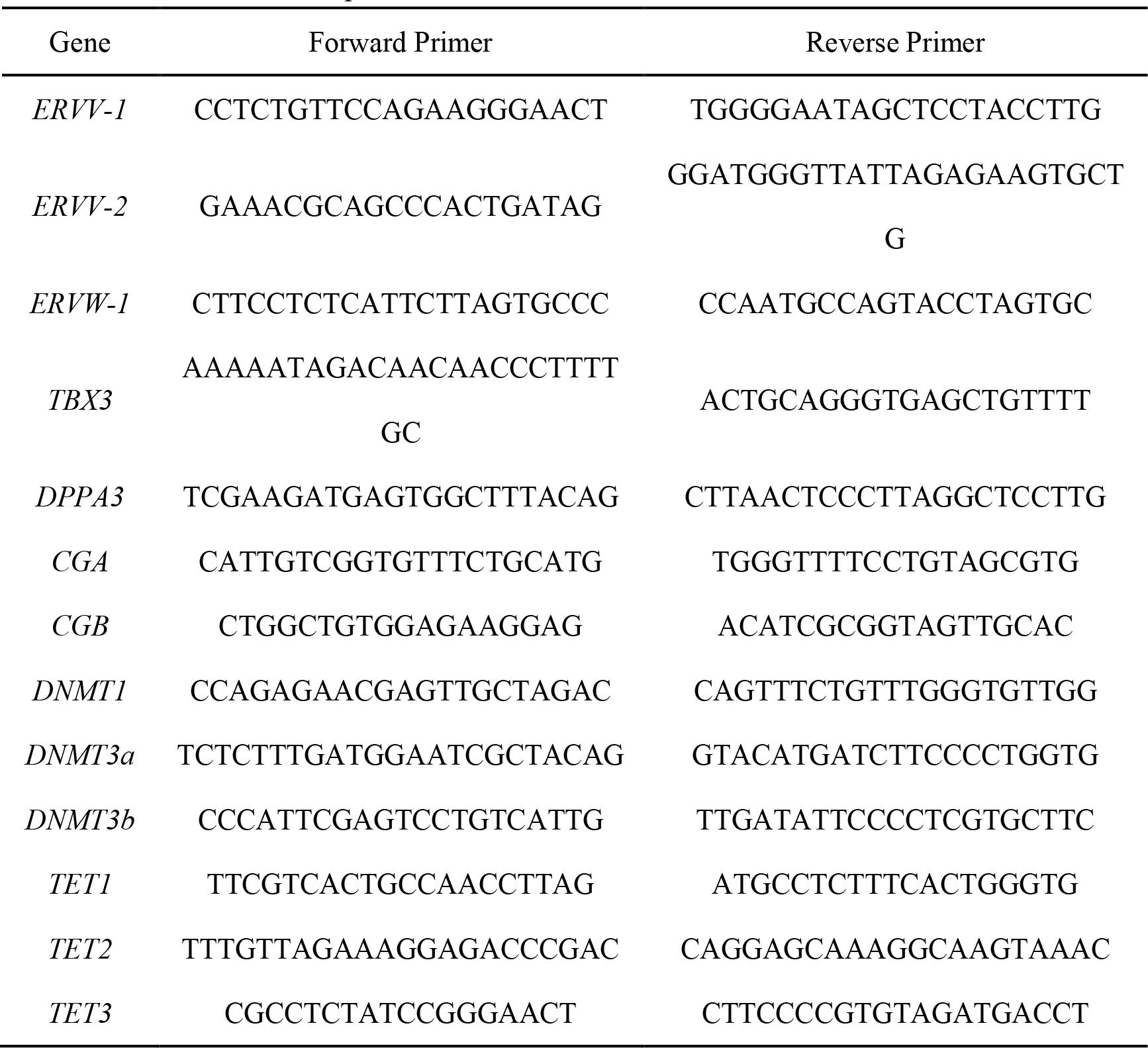
Primers used for qRT-PCR

